# Loss of developmental diapause as a prerequisite for social evolution in bees

**DOI:** 10.1101/649897

**Authors:** Priscila Karla Ferreira Santos, Maria Cristina Arias, Karen M. Kapheim

**Author notes:** Corresponding authors: Priscila K. F. Santos; and Karen M. Kapheim.

## Abstract

Diapause is a physiological arrest of development ahead of adverse environmental conditions and is a critical phase of the life cycle of many insects. In bees, diapause has been reported in species from all seven taxonomic families. However, they exhibit a variety of diapause strategies. These different strategies are of particular interest since shifts in the phase of the insect life cycle in which diapause occurs has been hypothesized to promote the evolution of sociality. Here we provide a comprehensive evaluation of this hypothesis with phylogenetic analysis and ancestral state reconstruction of the ecological and evolutionary factors associated with diapause phase. We find that social lifestyle, latitude, and voltinism are significant predictors of the life stage in which diapause occurs. Ancestral state reconstruction revealed that the most recent common ancestor of all bees likely exhibited developmental diapause and shifts to adult or reproductive diapause have occurred in the ancestors of lineages in which social behavior has evolved. These results provide fresh insight regarding the role of diapause as a prerequisite for the evolution of sociality in bees.

## Introduction

Diapause is a critical phase of the life cycle of many insects, and likely contributed to the ecological success of this highly diverse group of animals [1]. Many terms have been used to describe this phase of dormancy in insects, including diapause, adult diapause, reproductive diapause, hibernation, adult-wintering and overwintering. A defining feature of all these terms is an arrest in development or activity that is hormonally programmed in advance of environmental adversities such as harsh winter, dry seasons, or food restriction [1,2]. Diapause may occur at any stage of life: egg, larval, pupal or adult [1,3], and metabolic suppression varies from a decrease in activity (diapause in adult phase) to complete developmental arrest (diapause during development) [4]. Diapause may also be obligatory or facultative. Most obligatory diapausers live at high latitudes and produce only one generation per year. Conversely, in warmer regions, there are multiple active generations before winter and only one will pass through diapause (facultative diapause), and in tropical regions without extreme seasonal variation in resources, many insects forego diapause [5].

Bees are a diverse group of holometabolous insects in the Order Hymenoptera encompassing more than 20,000 species in seven taxonomic families [6]. Diapause has been reported in species of each family, with a great deal of variation in strategies. Although diapause in bees may occur in any developmental phase, the vast majority of species diapause during the last larval instar, immediately prior to metamorphosis, called the prepupal phase [7,8] or in the adult phase after mating and before the foundation of a new nest [9,10]. Also, a considerable number of species diapause as reproductively active adults, which is also known as reproductive diapause [11,12]. Some bee species are active year round and do not diapause [13,14].

The diversity of diapause strategies among bees is of particular interest since the developmental timing of this phenomenon plays a central role in key hypotheses for the evolution of sociality in Hymenoptera. Bees are an excellent system to explore these hypotheses, because sociality has evolved at least four times in two different families (Apidae – Apinae and Xylocopinae; Halictidae – Halictinae [Halictini, Augochlorini]), and species in each of these groups exhibit a great deal of variation in social lifestyles [15–18]. A feature common to each of these independent origins of sociality is cooperative nest sharing among adult females. In eusocial species, cooperating females are mothers and daughters who occupy reproductive castes, with non-reproductive daughters (i.e., workers) foregoing direct reproduction to help their mothers (i.e., queens) raise their siblings [19].

It has long been recognized that the haplodiploid mating system of the Hymenoptera promotes the evolution of eusociality, through its effects on sex allocation. As a result of males and females developing from unfertilized and fertilized eggs, respectively, females are more closely related to their sisters than they are to their own offspring [20,21]. This means that eusociality is expected to evolve from a solitary ancestor when some nests invest more in producing females, and other nests bias their reproductive investment toward males [22]. This split sex ratio provides an inclusive fitness benefit to helpers in female-biased nests, while maintaining a Fisherian sex ratio at the population level. Evidence for split sex ratios have been found between eusocial and semisocial nests [23] and eusocial and solitary nests [24] in facultatively social halictid bees.

Subsequent models for the origins of sociality have recognized that split sex ratios can also arise from temporally segregated patterns of sex allocation favored by shifts in the timing of diapause [25–27]. These models assume the production of at least two partially-overlapping generations per year (i.e., partial bivoltinism), and find that a female-biased sex ratio is favored in the second (summer) generation when females overwinter as adults after mating, but a male-biased sex ratio is favored when both females and males overwinter as larvae (Seger 1983). As such, eusociality is expected to evolve more readily in species that diapause as adults (Seger 1983, Quiñones and Pen 2017).

Diapause strategy has also been implicated in the mechanisms underpinning the evolution of sociality. Hunt and Amdam (2005) proposed the bivoltine ground plan (or diapause ground plan) hypothesis to explain the evolution of sociality in *Polistes* wasps. They hypothesized that physiological and behavioral differences between the non-diapausing (spring) and diapausing (summer) generations of an ancestral partially bivoltine solitary wasp could be co-opted to produce the worker and queen phenotypes, respectively [26,28]. This hypothesis also predicts that an ancestral shift from diapause as larvae to diapause as adults would have preceded the evolution of sociality.

Despite the prominence of diapause in models predicting the conditions that led to the evolution of sociality, there has not been a comprehensive evaluation of how diapause strategy corresponds to evolutionary transitions in sociality. If diapause of adult mated females is a necessary pre-adaptation for sociality to evolve, then shifts in the developmental timing of diapause should coincide with independent origins of sociality among the Hymenoptera. We tested this prediction by performing a phylogenetic analysis of diapause type as a function of social organization and other ecological traits among bees. We also used ancestral state reconstruction to characterize evolutionary transitions in diapause strategy within bee lineages in which social behavior has evolved. We found that diapause type is significantly correlated with social lifestyle, latitude, and voltinism and that shifts from prepupae to adult or reproductive diapause are likely to have preceded all independent origins of sociality in bees. These results provide support for the hypothesis that loss of developmental diapause is a prerequisite for the evolution of sociality in bees.

## Material and Methods

We reviewed the literature to collect information about diapause for individual bee species (Supplementary Table S1). We assigned a diapause type to each species according to the following definitions: development (during pre-imaginal stage, i.e. larva or prepupa phases), adult (hibernation before or after mating among adults), reproductive (a temporary halt in egg-laying among adults) [11,12], no diapause (no disruption in activity throughout the year) and plastic (species capable of more than one diapause type within the same population). We also classified species as solitary or social. Species with independent reproduction, but which nest in aggregations or share nests, but do not have reproductive castes (i.e., communal species) were considered solitary. Species in which females share nests and exhibit some kind of reproductive division of labor were considered social (e.g., sub-social, semi-social, primitively eusocial, and advanced eusocial) [29]). Facultatively social species were included as two different populations, with one being labeled solitary and the other social. This is because most facultatively social species (e.g., *Halictus rubicundus*) exhibit variation in social behavior at the population level, and this typically corresponds to differences in latitude and voltinism. We also considered ecological factors that may influence diapause type, including nesting patterns (i.e., in the ground or in cavities), and voltinism (univoltine or multivoltine, one or more than two generations per year, respectively). We recorded the latitude for each population.

### Correlation analyses

To identify whether diapause type is significantly correlated with sociality and ecological features, we analyzed 100 species for which we could obtain a complete data set (Supplementary Table S1). Facultatively social species or those with intrapopulation plasticity in diapause type were removed from these analyses.

We used phylogenetic generalized least square (PGLS) analysis to account for the effect of shared evolutionary history among the species [30]. We used the function gls in the R package nlme [31], assuming the Brownian motion model of evolution, to predict diapause type based on sociality, voltinism, latitude, nest type, and the interactions between sociality and voltinism, latitude and voltinism, and sociality and latitude (diapause type ~ sociality*voltinism + latitude*voltinism + sociality*latitude + nest). The model was fit to the data with maximum likelihood (ML). The extraction of the model coefficients, F-value, p-value and the comparison between the best models and the null model were performed using the functions coef and anova from stats [32] R package. The function stepAIC from MASS R package [33] was used to compare and identify the best model based on AIC values. The final dataset used in the correlations is available at Supplementary Material - Table S1 and the complete R output can be accessed at https://github.com/pkfsantos/Diapause_in_bees.

### Ancestral state reconstruction (ASR)

The tree for the ASR analysis was built using Mesquite v.3-6 [34], and the topology and branch lengths were added based on current molecular dated phylogenies. Most branch lengths are based on molecular distance between tribes or sub-families, but genus level branch lengths were applied when available. The references used to determine the topologies and branch lengths for each group are listed in the Supplementary Table S2. The branch length value for each species is available in the Supplementary Table S1.

The ancestral state reconstruction was run using the R packages phytools [35] and geiger [36]. We selected the best fitting model of evolution (equal-rates model - ER, all-rates-different model - ARD or symmetrical model - SYM), based on the smallest AIC and the greatest log-likelihood value. The make.simmap function was used for ancestral reconstruction using the empirical Bayesian approach, model of evolution ARD, and estimated pi to estimate the prior distribution on the root node of the tree.

## Results

We collected information about diapause type for 157 populations (105 solitary and 52 social) of 152 species (101 solitary and 51 social) from the seven taxonomic families of bees. From those species, 75 (47,7%) diapause during pre-imaginal stages of development; 47 (29,9%) diapause as adults; 15 (9.6%) engage in reproductive diapause; 15 (9,6%) have no diapause; and 5 (3,2%) were plastic (diapause switching between development or adult phase) (Figure 1). Adult diapause is more common among social species (around 49% of the species). In contrast, only 17.8% of solitary species diapause as adults. This includes several species in the Megachilidae tribe Osmiini, especially from the genus *Osmia*, and two species from the Apidae tribe Anthophorini and the andrenid *Andrena vaga*. Strikingly, we could not find any record of social species that diapause during development, though *Exoneurella lawson*i can exhibits plasticity between adult and developmental diapause (Supplementary Table S1).

**Figure 1:**
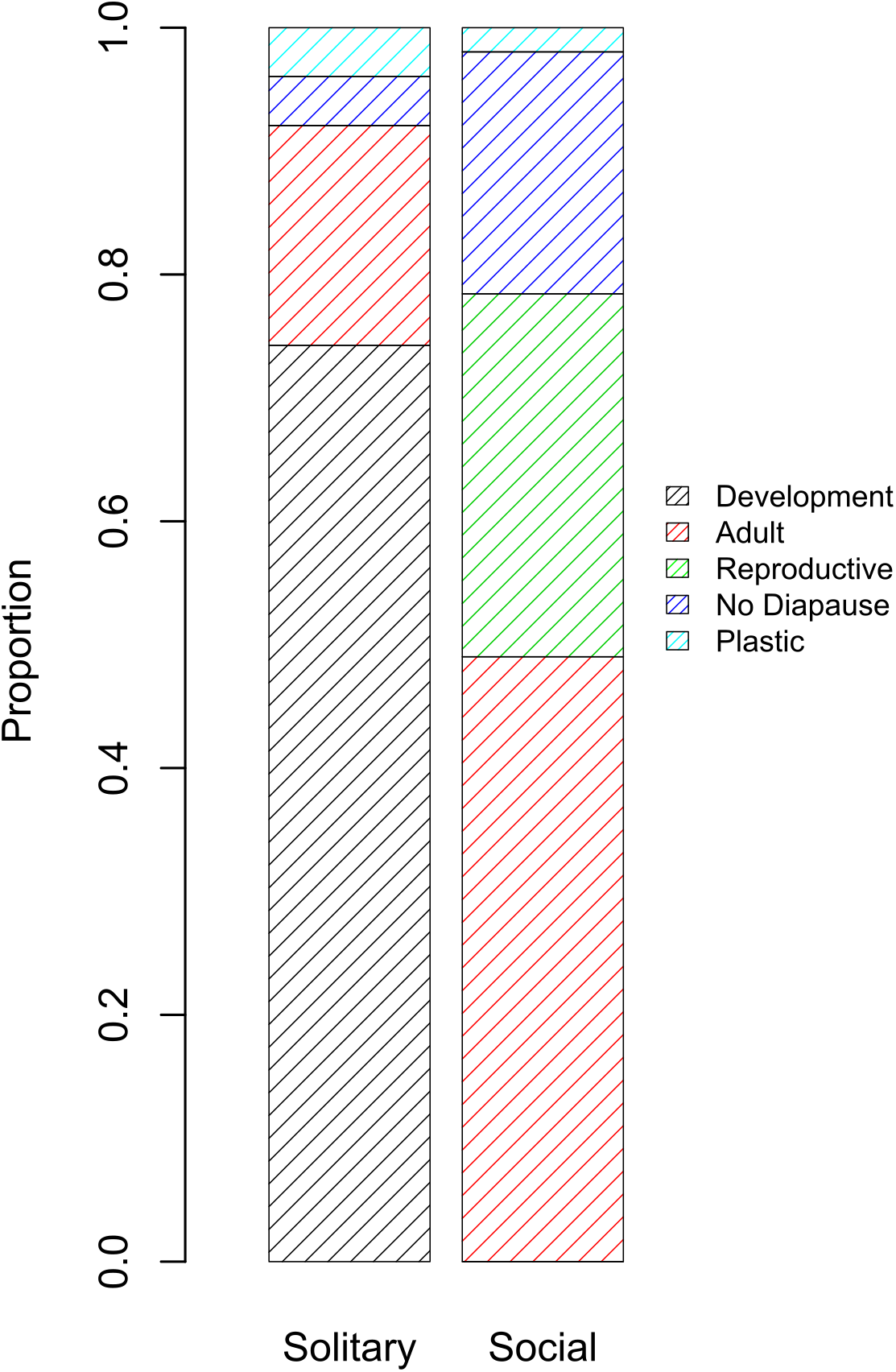
Proportion of species that enter in diapause in each category (development, adult, reproductive, no diapause or plastic). The species were divided in two classes according to their social lifestyle.

Our phylogenetic analysis revealed that diapause type is associated with variation in social organization, as well as ecological factors. The best fitting model from the PGLS analysis included sociality, voltinism, latitude, the interaction between voltinism and sociality, and the interaction between voltinism and latitude. This model was significantly better than the null model (logLik = −89.89, d.f. = 8, AIC difference > 10, p < 0.0001). Significant predictors of diapause type included sociality (F_1,93_ = 9.6, p = 0.0026), voltinism (F_1,93_ = 31.3, p < 0.0001), latitude (F_1,93_ = 53.4, p < 0.0001) and the interaction between voltinism and latitude (F_1,93_ = 26.9, p < 0.0001). Developmental diapause is only present in solitary species, while reproductive diapause only in social ones. Species with adult diapause are more frequent at high latitudes and those that do not diapause are more frequent at low latitudes. Univoltine species at high latitudes usually diapause during development or adult phase, while multivoltine species at low latitudes do not diapause or have a reproductive diapause.

Ancestral state reconstruction suggested that the ancestor of all bees diapaused during development (Figure 2). A shift from development to adult diapause is predicted to have occurred in the common ancestor of two different lineages, each of which has hosted an independent origin of sociality: once in Halictidae (Augochlorini+Halictini ancestors) and once in Apidae (Xylocopinae) (Figure 2). The other shift to adult diapause is predicted to have occurred in the ancestor of the Bombini, but in this case from an ancestor with reproductive diapause.

**Figure 2:**
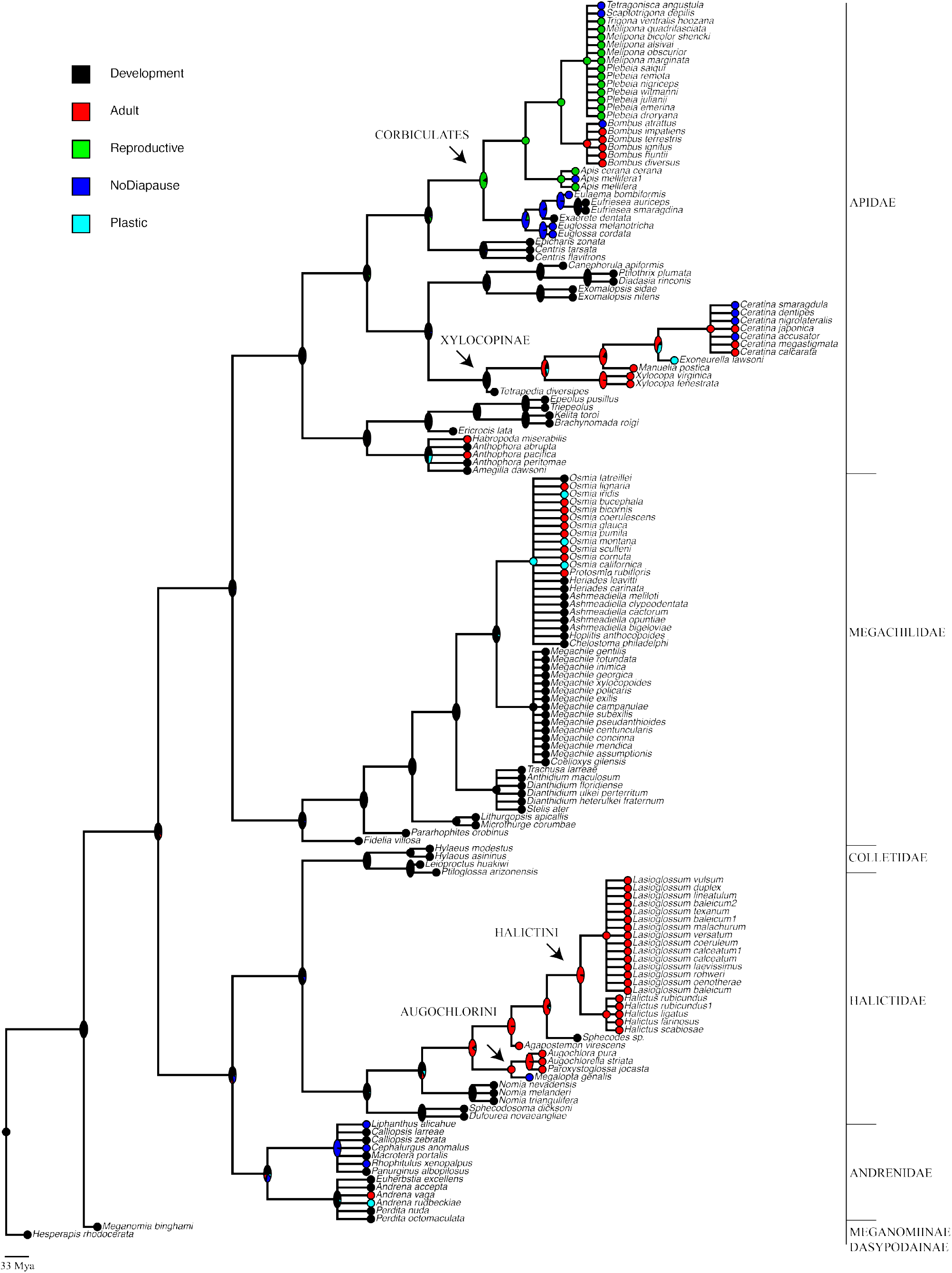
Ancestral reconstruction of diapause type in bees. Different colors represent the five diapause strategies: black - development, red - adult, green – reproductive, blue - no diapause and cyan – plastic (individuals may present developmental or adult diapause in the same population). The arrows are pointing to the groups which sociality evolved. The family and sub-families were classified accordingly with Cardinal and Danforth (2013).

The corbiculate bees (Apini, Bombini, Euglossini and Meliponini) are the only social lineage with an ancestor that has no predicted probability of adult diapause. Nonetheless, ancestral state reconstruction suggests that the ancestor of this group was likely to have shifted from developmental to reproductive diapause, which also occurs among adults. However, this result may be biased due to the large number of stingless bees in our dataset, most of which are tropical and exhibit reproductive diapause (Figure 2).

## Discussion

### Diapause strategy is highly variable among bees

Diapause in bees is extremely variable and, as in other groups of insects, may occur in any stage of life. Most bee species diapause as prepupae or adults, however there are a few peculiarities. For example, a *Sphecodes* sp., parasite of *Perdita nuda* diapause as postfed-predefecated larvae to mimic the host larva in its feeding and overwintering strategy [37]. This differs from other bees that diapause after defecation and from other closely related Halictidae species that diapause as adults (Figure 2).

Diapause is usually described as restricted to one stage of the life cycle for any given species [1,3]. In a recent review of diapause in insects, it was stated that there is no known case of diapause occurring in more than one stage in the same life cycle [38]. However, bees provide exception to this rule. For example, some *Osmia* species can diapause as either adults or prepupae, depending on whether they initiate a one or two year development strategy [39,40]. Individuals that develop in one year overwinter as adults, and individuals that extend development into a second year will undergo two periods of diapause, as prepupae in the first winter and as adults over the second winter [40]. Neff and Simpson (1997) found adults and prepupae of *Andrena rudbeckiae* overwintering in the same season and location [41]. Michener (1964) described *Exoneurella lawsoni* groups of adult and larvae of different ages overwintering in the same nest. Also, for *Centris tarsata* indirect evidence suggests that it presents both the adult and larval type diapause, depending on geographical location [42,43]. It is possible that this plasticity was also present in the ancestor of social lineages and provided the variation necessary for evolutionary shifts in diapause type that allowed for sociality to evolve.

The length of diapause is also variable among bees. Many species require an additional year or more to reach the maturity. *Osmia montana*, *O. californica,* and *O. iridis* complete development in either one or two years [40]. *Perdita nuda* and the parasite *Sphecodes* sp. may remain in diapause for up to 35 months [37]. Part of the population of *Euherbstia excellens* and some *Macrotera portalis* may take two or more years to complete the development [44,45]. Some *Chelostoma philadelphia* and *Diadasia rinconis* prepupae may spend an additional year in diapause [46,47], and for *Pararhophites orobinus* at least one prepupa is known to have remained in diapause for more than five years [48]. This plasticity in diapause length is suggested to be an adaptation to short growing seasons at high latitudes, which are insufficient to accomplish the development in a single season [5]. However, most species exhibiting this plasticity show intra-populational variation. As such, this is likely to be a bet-hedging mechanism common in populations with unpredictable variation in resource availability, ensuring that some individuals are likely to complete the life cycle when environmental conditions turn favorable [45,47].

### Diapause and the evolution of sociality

Theoretical models predict that eusociality is most likely to evolve in partially bivoltine species that pass unfavorable seasons as mated adult females [25–27]. This diapause strategy, along with other life history and ecological characteristics, promotes a female biased sex ratio in the summer generation, providing inclusive fitness benefits to helpers born in the spring generation [25,27]. The physiological, behavioral, and molecular mechanisms of diapause may also have been co-opted for the evolution of social castes [26,28]. Although these hypotheses share the premise that a departure from developmental diapause is a critical preadaptation for the evolution of sociality in Hymenoptera, diapause has never been comprehensively evaluated as a function of sociality. Our analysis provides multiple lines of support for the prediction that shifts in diapause phase facilitate the evolution of sociality in bees. We found that sociality is a significant predictor of diapause type in bees, in that all social bees that diapause do so as adults (adult or reproductive). Moreover, developmental diapause has been lost in the ancestor of all social lineages. Finally, we find that diapause type is significantly correlated with voltinism and latitude, other traits that have been postulated as important for the evolution of sociality [49,50].

Most social species diapause as adults or do not diapause, whereas most solitary species diapause during development. Even among those few solitary species that diapause as adults, there are important differences that set them apart from the social species, and potentially inhibit the evolution of eusociality. For example, species from the *Osmia* genus of solitary bees diapause as adults, but remain inside their pupal cocoon [51]. Diapause inside a cocoon may prevent interactions between mother and offspring in *Osmia*, which have important implications for social behavior in the adult phase in social and sub-social species [29,52–54]. The subsocial species *Xylocopa virginica* and *Ceratina calcarata* hibernate as adults in their natal nest prior to mating [55–57]. In contrast, species from the predominately eusocial genera *Lasioglossum*, *Halictus* and *Bombus* mate before passing the unfavorable period as adults [58–60]. This is an important distinction, because mating prior to diapause is predicted to facilitate the evolution of eusociality due to the effects it has on offspring sex ratio the following season [25,27].

Ancestral state reconstruction analysis suggested that the shifts from development to adult diapause correspond with the evolution of sociality in the same groups. Sociality has arisen in the family Halictidae two or three times, once in Augochlorini and either once or twice in Halictini [15,18]. We found that a shift from development to adult diapause is likely to have occurred once in the Augochlorini+Halictini ancestor. There have also been several reversals from a social to a solitary lifestyle in the Halictidae [61], and these species all diapause as adults. Sociality has also arisen in the subfamily Xylocopinae [17], and this corresponds to a shift from developmental to adult diapause.

One exception to this pattern is among the corbiculate bees, whose ancestor is predicted to have shifted from developmental to reproductive diapause. The corbiculates include the honey bees and stingless bees, which have the most advanced forms of eusociality among the bees, including perennial colonies and distinct morphological castes [62]. Quiñones and Pen (2017) demonstrate that the conditions that favor adult female diapause are not necessary once morphological castes evolve, due to feedback between helping behavior and sex allocation. Nonetheless, reproductive diapause is likely to also yield the conditions that favor a female-biased sex ratio, and thus the evolution of sociality, though this has not been specifically addressed in theory. Reproductive diapause occurs among adults, but the associated physiological changes are less intense than in adult diapause. Thus, the loss of developmental diapause may bring about the initial physiological change necessary for sociality to evolve.

## Conclusions

Diapause in bees is an extremely variable phenotype. It may occur in different phases of the life cycle and is variable in length. This diversity allowed a phylogenetic test of the role of diapause type in social evolution, revealing that the diapause type is significantly correlated to sociality, voltinism and latitude. Interestingly, developmental diapause does not occur in social species, and ancestral shifts from developmental to adult or reproductive diapause precede the evolution of social behavior. This suggest that the loss of the developmental diapause is an important preadaptation to sociality, and that the diapause ground plan proposed to have been co-opted for sociality in wasps may also apply to the evolution of sociality in bees.

## Acknowledgements

We would like to thank Dr Carlos Garófalo and Nicholas Saleh for information about *Centris* and *Euglossa* species, respectively; Tim Delory for the discussions; and Dr Klaus Hartfelder for providing valuable feedback on an earlier version of this manuscript.

